# Transcriptome Differences between Alternative Sex Determining Genotypes in the House Fly, *Musca domestica*

**DOI:** 10.1101/016774

**Authors:** Richard P. Meisel, Jeffrey G. Scott, Andrew G. Clark

## Abstract

Sex determination evolves rapidly, often because of turnover of the genes at the top of the pathway. The house fly, *Musca domestica*, has a multifactorial sex determination system, allowing us to identify the selective forces responsible for the evolutionary turnover of sex determination in action. There is a male determining factor, *M*, on the *Y* chromosome (*Y*^*M*^), which is probably the ancestral state. An *M* factor on the third chromosome (*III^M^*) has reached high frequencies in multiple populations across the world, but the evolutionary forces responsible for the invasion of *III^M^* are not resolved. To test if the *III*^*M*^ chromosome invaded because of sex-specific selection pressures, we used mRNA sequencing to determine if isogenic males that differ only in the presence of the *Y*^*M*^ or *III*^*M*^ chromosome have different gene expression profiles. We find that more genes are differentially expressed between *Y*^*M*^ and *III*^*M*^ males in testis than head, and that genes with male-biased expression are most likely to be differentially expressed between *Y*^*M*^ and *III*^*M*^ males. We additionally find that *III*^*M*^ males have a “masculinized” gene expression profile, suggesting that the *III*^*M*^ chromosome has accumulated an excess of male-beneficial alleles because of its male-limited transmission. These results are consistent with the hypothesis that sex-specific selection acts on alleles linked to the male-determining locus driving evolutionary turnover in the sex determination pathway.

## 1. Introduction

Sex determination (SD) is an essential developmental process responsible for sexually dimorphic phenotypes. It is therefore paradoxical that SD pathways are poorly conserved, with master SD (MSD) genes at the top of the pathway differing between closely related species and even variable within species (Bull, 1983; Wilkins, 1995; Pomiankowski et al., 2004; Beukeboom and Perrin, 2014). The hypotheses to explain the rapid evolution of SD pathways fall into three categories. First, SD evolution may be selectively neutral if MSD turnover is the result of mutational input without phenotypic or fitness consequences (van Doorn, 2014). Second, frequency dependent (sex-ratio) selection could favor a new MSD variant if one sex is below its equilibrium frequency (Eshel, 1975; Bull and Charnov, 1977; Bulmer and Bull, 1982; Werren and Beukeboom, 1998). Third, a new MSD locus can invade a population if the new MSD variant itself or genetically linked alleles confer a fitness benefit (Charlesworth and Charlesworth, 1980; Rice, 1986; Charlesworth, 1991; van Doorn and Kirkpatrick, 2007, 2010). Those fitness effects could be beneficial to both sexes (natural selection), increase the reproductive success of the sex determined by the new MSD variant (sexual selection), or be beneficial to the sex determined by the MSD variant and deleterious to the other sex (sexually antagonistic selection). Sexually antagonistic selection is predicted to be an especially important driver of MSD turnover because linkage to an MSD locus allows the sexually antagonistic allele to be inherited in a sex-limited manner, thereby resolving the inter-sexual conflict (Charlesworth and Charlesworth, 1980; van Doorn and Kirkpatrick, 2007; Roberts et al., 2009; van Doorn and Kirkpatrick, 2010).

The house fly, *Musca domestica*, is an ideal model for testing hypotheses about the evolution of SD because it has a multifactorial SD system, with male- and female-determining loci segregating in natural populations (Dübendorfer et al., 2002; Hamm et al., 2015). Most relevant to the work presented here is the the fact that the male-determining factor, *M*, can be located on the *Y* chromosome (*Y*^*M*^), any of the five autosomes (*A*^*M*^), and even the *X* chromosome (Hamm et al., 2015). It is unknown whether these *M*-factors are the same gene in different locations or different genes that have independently assumed the role of an MSD locus (Bopp, 2010). *Y*^*M*^ is a common arrangement (Hamm et al., 2015), and it is thought to be the ancestral state because it is the genotype found in close relatives of the house fly (Boyes et al., 1964; Boyes and Van Brink, 1965; Dübendorfer et al., 2002). *M* on the third chromosome (*III*^*M*^) is also common, but it is not clear what was responsible for the invasion of the *III*^*M*^ chromosome (Hamm et al., 2015). Note that when the *M* factor arrived on chromosome *III*, this entire chromosome essentially assumed *Y*-like properties, including male-biased transmission and reduced recombination (Hamm et al., 2015). However, the *III*^*M*^ chromosome is not a degenerated *Y* chromosome because *III*^*M*^ homozygotes are viable, fertile, and found in natural populations (Hamm et al., 2015). Identifying the selective forces responsible for the invasion of *III*^*M*^ will be a powerful test of the hypotheses to explain SD evolution.

Strong linkage to *A*^*M*^ is expected for alleles on the same autosome because recombination is low or non-existent in house fly males (Hiroyoshi, 1961; Hamm et al., 2015), but see Feldmeyer et al. (2010). It is possible that *A^M^* chromosomes invaded house fly populations because of selection on phenotypic effects of either the autosomal *M* loci themselves or alleles linked to *M*-factors (Franco et al., 1982; Tomita and Wada, 1989; Kozielska et al., 2006; Feldmeyer et al., 2008). *M* variants are known to have subtle phenotypic effects, which include differential splicing and expression of SD pathway genes between *Y*^*M*^ and *A*^*M*^ males (Schmidt et al., 1997; Hediger et al., 2004; Siegenthaler et al., 2009). In addition, *A*^*M*^ chromosomes form stable latitudinal clines on multiple continents (Franco et al., 1982; Tomita and Wada, 1989; Hamm et al., 2005; Kozielska et al., 2008), and seasonality in temperature is somewhat predictive of their distribution (Feldmeyer et al., 2008). Furthermore, in laboratory experiments, *III*^*M*^ males out-competed *Y*^*M*^ males for female mates; the *III*^*M*^ chromosome increased in frequency over generations in population cages; and *III*^*M*^ males had higher rates of emergence from pupae than *Y*^*M*^ males (Hamm et al., 2009). The most specific phenotype that has been linked to *A*^*M*^ is insecticide resistance (Kerr, 1960, 1961, 1970; Denholm et al., 1983; Kence and Kence, 1992), but insecticide resistance alone cannot entirely explain the invasion of *A*^*M*^ chromosomes (Shono and Scott, 1990; Hamm et al., 2005). These results all support the hypothesis that natural, sexual, or sexually antagonistic selection on *M* variants or linked alleles drove the invasion of *A*^*M*^ chromosomes.

To test whether sex-specific selection pressures could be responsible for the invasion of the *III*^*M*^ chromosome, we used high-throughput mRNA sequencing (mRNA-Seq) to compare gene expression profiles between nearly isogenic *Y*^*M*^ and *III*^*M*^ males that only differ in their *M*-bearing chromosome. These contrasts are essentially a comparison between flies with the ancestral *Y* chromosome (*Y*^*M*^) and individuals with a recently evolved “neo-Y” (*III*^*M*^). The gene expression differences we detected were the result of both differentiation of *cis* regulatory regions between the *III*^*M*^ and “standard” third chromosome and *trans* effects of the *III*^*M*^ and/or *Y*^*M*^ chromosome(s) on expression throughout the genome. We found that genes responsible for male phenotypes are more likely to be differentially expressed between *Y*^*M*^ and *III*^*M*^ males, suggesting that *Y*^*M*^ and *III*^*M*^ males have phenotypic differences that would be differentially affected by male-specific selection pressures. We also found that *III*^*M*^ males have a “masculinized” gene expression profile. These results support the hypothesis that sexual or sexually antagonistic selection drives evolutionary turnover at the top of SD pathways.

## 2. Materials and Methods

### 2.1. Strains

We compared gene expression between two nearly isogenic house fly strains that differ only in the chromosome carrying *M*. The first, Cornell susceptible (CS), is an inbred, lab adapted strain with *XX* males that are heterozygous for a *III*^*M*^ chromosome and a standard third chromosome that lacks an *M* factor (*X/X*; *III*^*M*^ /*III*^*CS*^) (Scott et al., 1996; Hamm et al., 2005) (Figure 1A). CS females are *XX* and homozygous for the standard third chromosome (*X/X*; *III*^*CS*^/*III*^*CS*^). We created a strain with *Y*^*M*^ males that has the *X* chromosome and all standard autosomes from the CS strain. To do so, we used a backcrossing approach to move the *Y* chromosome from the genome strain (aabys) onto the CS background (Figure 1B), creating the strain CS-aabys-Y (CSaY). CSaY males are *XY* and homozygous for the standard CS third chromosome (*X/Y*; *III*^*CS*^/*III*^*CS*^). The aabys strain has a recessive phenotypic marker on each of the five autosomes (Wagoner, 1967; Tomita and Wada, 1989). To confirm that the aabys autosomes had been purged from the CSaY genome, we crossed CSaY flies to aabys and observed only wild-type progeny. CS and CSaY males are nearly isogenic, differing only in that CS males are *XX* and heterozygous for the *III*^*M*^ and standard *III*^*CS*^ chromosomes, and CSaY males are *XY* and homozygous for the standard *III*^*CS*^ chromosome (Figure 1). Females are genetically identical between strains.

**Figure 1:**
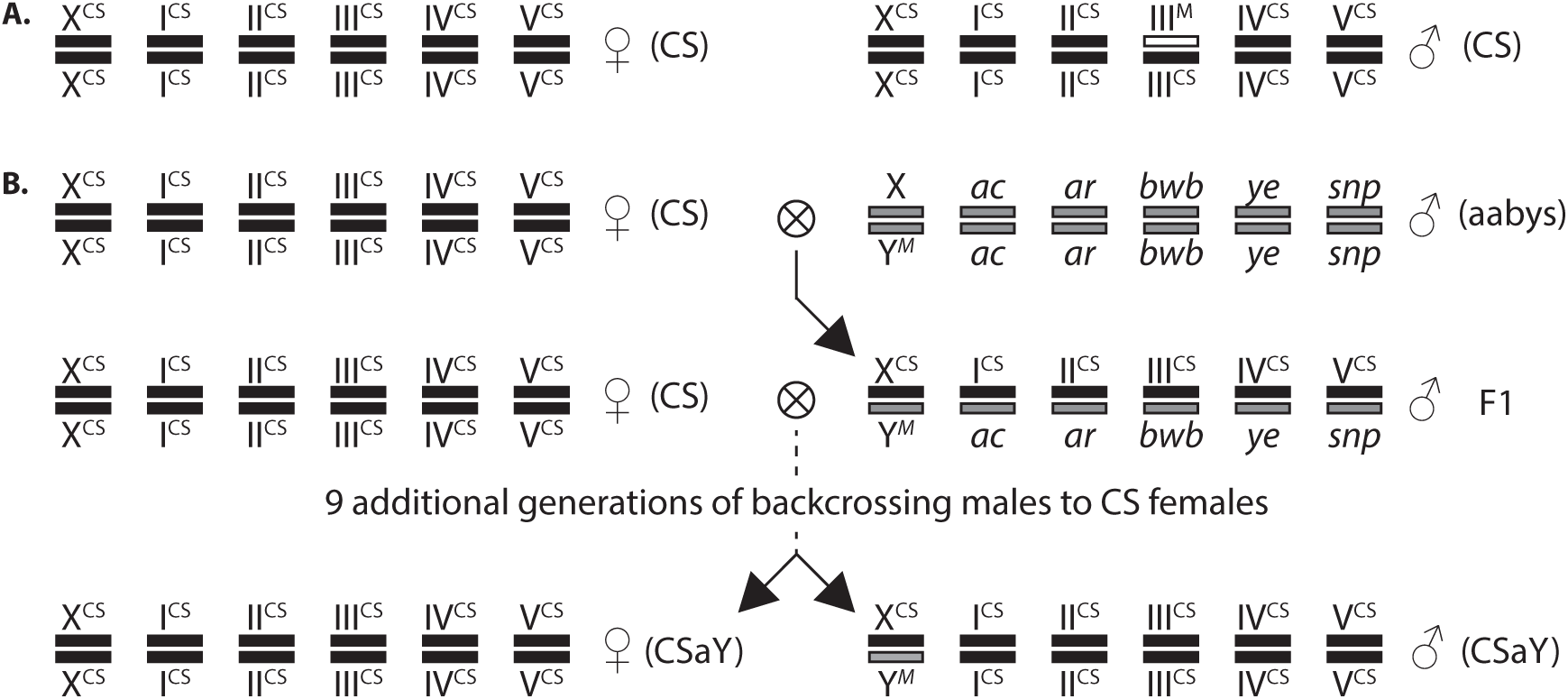
Genotypes of flies. **A.** Genotypes of CS males and females. **B.** Crossing scheme used to generate the CSaY (Y^M^) strain. Each pair of parallel rectangles represent homologous chromosomes; there is one pair of sex chromosomes (X and Y) and five autosomes. Chromosomes of CS origin are black and indicated by “CS”, except for the III^M^ chromosome which is white. Chromosomes of aabys origin are gray, and the aabys strain has a recessive phenotypic marker on each autosome. Females from the CS strain were crossed to aabys males, and the male progeny were backcrossed to CS females for 10 generations to create the CSaY strain. Because there is no recombination in XY males, CS and CSaY are isogenic except that the CSaY males have a Y chromosome and CS males have a III^M^ chromosome.

We are confident that the strains are isogenic except for the *M*-bearing chromosome because there is very little evidence for recombination in male house flies with an *XY* genotype (Hiroyoshi, 1961; Hamm et al., 2015). However, if there were minimal recombination between the CS and aabys chromosomes in our crossing scheme, the majority of autosomal alleles in the CSaY strain would still have originated from the CS genotype, with very little contribution from aabys autosomes.

### 2.2. Samples and mRNA-Seq

CS and CSaY flies were kept at 25°C with a 12:12 hour light:dark cycle. Larvae were reared in media made with 1.8 L water, 500 g calf manna (Manna Pro, St. Louis, MO), 120 g bird and reptile litter wood chips (Northeastern Products, Warrensburg, NY), 60 g dry active baker’s yeast (MP Biomedical Solon, OH), and 1.21 kg wheat bran (Cargill Animal Nutrition, Minneapolis, MN), as described previously (Hamm et al., 2009).

We sampled two types of tissue from CS and CSaY males and females: head and gonad. All dissections were performed on living, non-anesthetized 4–6 day old unmated adult flies. Heads were separated from males and females, homogenized in TRIzol reagent (Life Technologies) using a motorized grinder, and RNA was extracted on QIAGEN RNeasy columns following the manufacturer’s instructions including a genomic DNA (gDNA) elimination step. Testes were dissected from males, and ovaries were dissected from females in Ringer’s solution (182 mM KCl, 46 mM NaCl, 3 mM CaCl_2_, 10 mM Tris-Cl in ddH_2_O). Ovary and testis samples were dissolved in TRIzol and RNA was extracted on QIAGEN RNeasy columns with gDNA elimination. Three biological replicates of CS (*III*^*M*^) male heads, CSaY (*Y*^*M*^) male heads, CS testes, and CSaY testes were collected; one sample was collected for each of the four female dissections (CS head, CSaY head, CS ovary, and CSaY ovary).

Barcoded mRNA-Seq libraries were prepared using the Illumina TruSeq kit following the manufacturer’s instructions. Briefly, mRNA was purified using oligo-dT magnetic beads, cDNA was synthesized using random hexamer primers, and sequencing libraries were constructed using the cDNA. The samples were run on 2 lanes of an Illumina HiSeq2500 at the Cornell Medical School Genomics Resources Core Facility. One lane had the eight head samples, and the other lane had the eight gonad (testis and ovary) samples. We generated 101 base pair single-end reads, and the sequencing reads were processed using Casava 1.8.2.

### 2.3. mRNA-Seq data analysis

Illumina mRNA-Seq reads were aligned to house fly genome assembly v2.0.2 and annotation release 100 (Scott et al., 2014) using TopHat2 v2.0.8b (Kim et al., 2013) and Bowtie v2.1.0.0 (Langmead et al., 2009) with the default parameters. We tested for differential expression between males and females and between *Y*^*M*^ and *III*^*M*^ males using the Cuffdiff program in the Cufflinks v2.2.1 package (Trapnell et al., 2013) with the default parameters, including geometric normalization. We used a false discovery rate (FDR) corrected *P*-value of 0.05 to identify genes that are differentially expressed (Benjamini and Hochberg, 1995). Genes were considered not differentially expressed if Cuffdiff returned an “OK” value for the test status (at least 10 reads aligned to the transcript, and data were sufficient for testing for differential expression) but the expression levels were not significantly different. Genes without an “OK” value were not included in downstream analyses. We repeated this analysis by also requiring a two-fold difference in expression to call genes as differentially expressed. In comparisons between male and female expression levels, we treated all 6 male samples as biological replicates and did the same for both female samples. We also repeated the analysis using only 2 replicates of each sample to control for sample-size effects.

We used expression level estimates from Cuffdiff2 (Fragments Per Kilobase of transcript per Million mapped reads, FPKM) to calculate correlations of expression levels between our experimental samples (Figure S1). Only genes with an “OK” value for test status in Cuffdiff were included. The correlations between testis and ovary expression are lowest, which is expected because they are dramatically different tissues. The correlations between male and female head samples are substantially higher than between testis and ovary, but still lower than the correlations within sexes. The two ovary samples are more highly correlated than any of the pairwise comparisons between CS and CSaY testis samples (Figure S1), most likely because CS and CSaY females are genetically identical (Figure 1). All data analysis was performed in the R statistical programming environment (R Core Team, 2015).

### 2.4. Chromosomal assignments of house fly genes

The house fly genomic scaffolds have not formally been assigned to chromosomes, but homologies have been inferred between house fly chromosomes and the five major chromosome arms of *Drosophila*, also known as Muller elements A–E (Foster et al., 1981; Weller and Foster, 1993). Additionally, the house fly *X* chromosome is most likely homologous to the *Drosophila* dot chromosome (Muller element F, or *Drosophila melanogaster* chromosome 4) (Vicoso and Bachtrog, 2013, 2015). We therefore assigned house fly genes that are conserved as one-to-one orthologs with *D. melanogaster* (Scott et al., 2014) to house fly chromosomes based on the Muller element mapping of the *D. melanogaster* orthologs. For our analysis of gene families that are differentially expressed between *Y*^*M*^ and *III*^*M*^ males, we assigned house fly scaffolds to chromosomes based on the Muller element mapping of the majority of *D. melanogaster* orthologs on each scaffold.

### 2.5. Gene ontology analysis

We used the predicted *D. melanogaster* orthologs (Scott et al., 2014) to infer the functions of house fly genes. Gene ontology (GO) annotations of house fly genes were determined using Blast2GO (Conesa et al., 2005; Götz et al., 2008) as described previously (Scott et al., 2014). We then used Blast2GO to identify GO classes that are enriched amongst differentially expressed genes relative to non-differentially expressed genes using an FDR-corrected Fisher’s exact test (Benjamini and Hochberg, 1995).

### 2.6. qPCR validation of differentially expressed genes

We used quantitative PCR (qPCR) on cDNA to validate the differential expression of four genes between *Y*^*M*^ and *III*^*M*^ testes. Dissections of testes were performed as described above on three batches of *Y*^*M*^ and *III*^*M*^ males each (i.e., 3 biological replicates of each strain). RNA was extracted from testes using TRIzol homogenization followed by purification using Directzol RNA MiniPrep columns (Zymo Research). We synthesized cDNA using M-MLV reverse transcriptase (Promega) with oligo-dT primers following the manufacturer’s instructions. We designed PCR primers to amplify a 71–110 bp product at the 3’ end of each transcript, with one primer on either side of the 3’-most annotated intron when possible (Table S1). We also designed primers to amplify one transcript that was not differentially expressed between *Y*^*M*^ and *III*^*M*^ males or between males and females in either gonad or head in our mRNA-Seq analysis (XM 005187313). We tested our primer pairs with PCR using testis cDNA as a template to validate that they amplify a single product.

We then performed qPCR on 3 technical replicates of 5 serial dilutions of 1/5 each using a 60°C annealing temperature on an Applied Biosystems StepOnePlus Real-Time PCR System with Power SYBR Green PCR Master Mix (Life Technologies) following the manufacturer’s instructions. We assigned a threshold cycle (C_T_) to the qPCR curves, and we validated that the primer pairs gave a linear relationship between C_T_ and –*log*_10_ concentration. We next used the same reagents and equipment to perform qPCR using each primer pair on 3 technical replicates of the 3 biological replicates from both strains (18 samples total), with the samples interspersed on a 96-well plate to avoid batch effects. In addition to the 18 samples, each qPCR plate contained 3 technical replicates of a 5-step 1/5 serial dilution (15 samples). Those 33 samples were amplified by qPCR using primers for a gene that was detected as differentially expressed using mRNA-Seq and the 33 samples were also amplified with primers in our control gene (XM 005187313), for a total of 66 reactions on a single plate. From each plate, we constructed a standard curve for each primer pair using the relationship between C_T_ and –*log*_10_ concentration from our serial dilutions, and we used the slope of these lines to estimate the initial concentration of our template cDNA in each sample. We divided the initial concentration for our experimental gene by the estimated concentration for the control gene to determine a relative concentration for the experimental gene for each of the technical replicates.

To test for differential expression between *Y*^*M*^ and *III*^*M*^ samples, we first constructed a linear model with replicates nested within strains predicting the relative concentration of the gene in the R statistical programming environment (R Core Team, 2015). We then used Tukey’s Honest Significant Differences method to perform an analysis of variance to determine if there is a significant effect of strain on expression level.

## 3. Results

### 3.1. Genes on the house fly third chromosome are more likely to be differentially expressed between Y^M^ and III^M^ males than genes on other autosomes

We used mRNA-Seq to compare gene expression in heads and gonads of house fly males and females of a *Y*^*M*^ strain (CSaY) and a *III*^*M*^ strain (CS). Males of the *III*^*M*^ strain are *XX* and heterozygous for the *III*^*M*^ chromosome and a standard third chromosome without *M* (Figure 1A). Males of the *Y*^*M*^ strain are *XY* (with the same *X* as the *III*^*M*^ strain) and homozygous for the standard third chromosome found in the *III*^*M*^ strain (Figure 1B). The rest of the genome is isogenic, and females of the two strains are genetically identical (Figure 1).

We detected 873 and 1338 genes that are differentially expressed between *Y*^*M*^ and *III*^*M*^ males in heads or testes, respectively, at an FDR-corrected *P*-value of 0.05 (Table 1, Figure S2, Supplementary Data). Genes on the house fly third chromosome are more likely than genes on other autosomes to be differentially expressed between *Y*^*M*^ and *III*^*M*^ males (Figure 2A). Approximately 30% of the differentially expressed genes are predicted to be on the third chromosome, which is greater than the fraction assigned to any of the other four autosomes (14.8–20.6%). There is a slight, but significant, signal of higher expression from the third chromosome in *Y*^*M*^ males when compared to *III*^*M*^ males (Figure 2B), and a significant excess of third chromosome genes is up-regulated in *Y*^*M*^ males (Figure S3). However, a significant excess of third chromosome genes is also up-regulated in *III*^*M*^ males (Figure S3), suggesting that the differential expression of third chromosome genes between *Y*^*M*^ and *III*^*M*^ males is not the result of degeneration of the *III*^*M*^ chromosome.

**Table 1:** Differential expression between strains and sexes. Counts of the number of genes that are differentially expressed (# diff), tested and non-differentially expressed (#non-diff), and total genes tested (# genes), as well as the frequency of genes that are differentially expressed (freq diff). For the # diff columns, the number of genes up-regulated in each of the two samples being compared are presented.

| Tissue | comparison | # diff |  | # non-diff | # genes | freq diff |
| --- | --- | --- | --- | --- | --- | --- |
| | | $Y^M$ | $III^M$ | | | |
| male head | $Y^M$ vs. $III^M$ | 320 | 553 | 8568 | 9441 | 0.092 |
| testis | $Y^M$ vs. $III^M$ | 782 | 556 | 8937 | 10275 | 0.130 |
|  |  | male | female |  |  |  |
| head | male vs. female | 205 | 1087 | 8387 | 9679 | 0.133 |
| gonad | testis vs. ovary | 3426 | 4369 | 2791 | 10586 | 0.736 |

**Figure 2:**
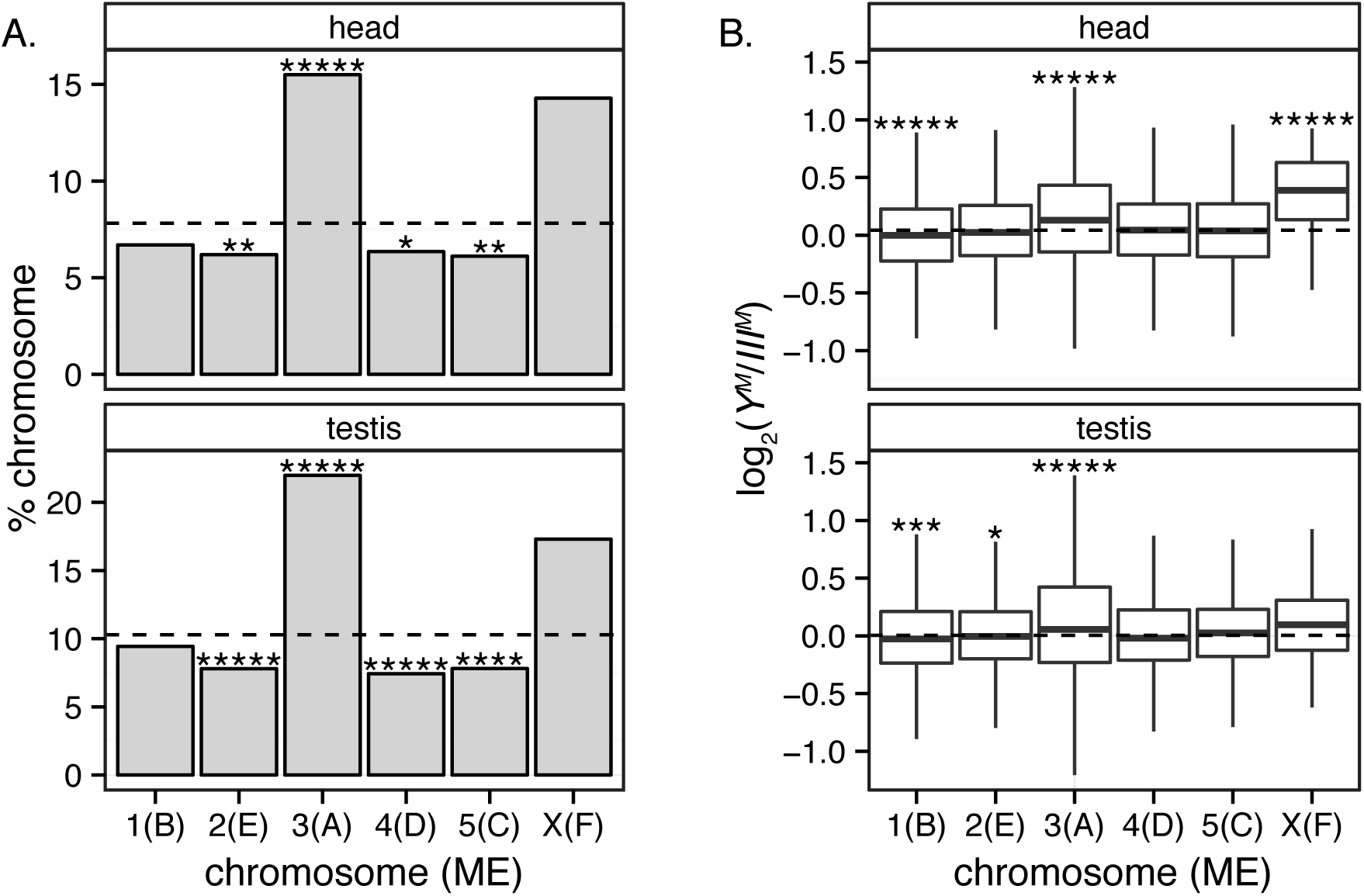
Chromosomal mapping and differential expression between Y^M^ and III^M^ males. **A.** Bar graphs indicate the percent of genes on each chromosome (Drosophila Muller element in parentheses) that are differentially expressed between Y^M^ and III^M^ male heads (top) or testes (bottom). The dashed line indicates the percentage of genes that are differentially expressed across all chromosomes. Asterisks indicate *P*-values from Fisher’s exact test comparing the number of differentially expressed genes with the number not differentially expressed on a chromosome and summed across all other chromosomes (*P < 0.05, **P < 0.005, ****P < 0.00005, *****P < 0.000005). **B.** Box plots show the relative expression levels of genes in Y^M^ and III^M^ males on each chromosome. Expression level was measured in head (top) and testis (bottom). The dashed line indicates the average *log_2_* expression ratio across all genes. Asterisks indicate *P*-values from a Mann-Whitney test comparing the expression ratio between genes on one chromosome versus the rest of the genome (*P < 0.05, **P < 0.005, ***P < 0.0005, ****P < 0.00005, *****P < 0.000005).

*X*-linked genes also trend towards an excess that are differentially expressed between *Y*^*M*^ and *III*^*M*^ males (Figure 2A), but we do not have the power to detect statistically significant deviations from the expectation because only 56 *X*-linked genes are expressed in head and 52 *X*-linked genes are expressed in testis. Surprisingly, expression from the *X* chromosome is higher in *Y*^*M*^ (*XY*) than *III*^*M*^ (*XX*) male heads (Figure 2B), and a significant excess of *X*linked genes is up-regulated in *Y*^*M*^ heads relative to *III*^*M*^ heads (Figure S3). These results demonstrate that differential expression of *X*-linked genes between *Y*^*M*^ and *III*^*M*^ males is not the result of a dosage deficiency in hemizygous males. In addition, these patterns suggest that either a dosage compensation mechanism provides greater than two-fold upregulation of the *X* chromosome in *XY* males or *trans* effects of the *Y* chromosome lead to up-regulation of *X*-linked expression.

Our chromosomal assignments of house fly genes are almost certainly less than perfect because of gene movement between chromosomes since the divergence between *D. melanogaster* the *M. domestica* lineages. However, errors in chromosomal assignments should obstruct the signal of elevated expression divergence on the third and *X* chromosomes, making our results conservative.

### 3.2. More differential expression between Y^M^ and III^M^ males in testis than in head, but a common set of genes co-regulated in both tissues

A higher fraction of genes is differentially expressed in testes between *Y^M^* and *III*^*M*^ males than in heads (Table 1; *P <* 10^−16^ in Fisher’s exact test, FET), suggesting that genes involved in male fertility phenotypes are more affected by the *M*-bearing chromosome. When we restricted the analysis to only genes expressed in both heads and gonads, we still observe more genes differentially expressed in testes than heads between *Y^M^* and *III*^*M*^ males (Table S2). When we used a two-fold cutoff in addition to an FDR-corrected *P <* 0.05 cutoff, the number of genes differentially expressed in head and testis between *Y*^*M*^ and *III*^*M*^ males goes down to 373 and 558, respectively. However, there is still a higher fraction of genes differentially expressed in testis than head (*P*_*FET*_ < 10^−5^)

Genes that are differentially expressed between males and females are said to have “sex-biased” expression (Ellegren and Parsch, 2007). The fraction of genes differentially expressed between the testes of *Y*^*M*^ and *III*^*M*^ males is nearly as large as the fraction with sex-biased expression in head (Table 1). We have a different number of replicates for male samples (three *Y*^*M*^ and three *III*^*M*^, for a total of six male replicates) than female samples (two), and so our power to detect differential expression may differ between the inter-strain (*Y^M^ vs. III^M^*) and inter-sex comparisons. To control for sample size effects, we repeated our analysis using only two replicates of each male strain in the inter-strain comparison and two male replicates (one from each strain) in the inter-sex comparison. With only two replicates of each sample we confirmed that more genes are differentially expressed between *Y*^*M*^ and *III*^*M*^ testes than heads, and we found that more genes are differentially expressed between *Y*^*M*^ and *III*^*M*^ males in either tissue than have sex-biased expression in head (Table S3). This result confirms that the inter-strain expression differences are of a similar or greater magnitude than the amount of sex-biased expression in head.

If the probability that a gene is differentially expressed between *Y*^*M*^ and *III*^*M*^ male heads is independent of the probability that the gene is differentially expressed in testes, we expect *<*1% of genes to be differentially expressed in both head and testis. We find that 176 genes (2.12%) are differentially expressed between *Y*^*M*^ and *III*^*M*^ males in both head and testis when using an FDR-corrected *P*-value to test for differential expression, which is significantly greater than the expectation (*P*_*FET*_ < 10^−25^). In contrast, there is not a significant excess of genes with sex-biased expression in both head and gonad—we expect 9.41% of genes to have sex-biased expression in both head and gonad (Table S2), and we observe that 809 genes (9.27%) are sex-biased in both tissue samples (*P*_*FET*_ = 0.655). We obtain qualitatively similar results when using a two-fold cutoff in addition to an FDRcorrected *P*-value threshold of 0.05 to test for differential expression: there is a four-fold excess of genes that are differentially expressed in both head and testis between *Y*^*M*^ and *III*^*M*^ males (P_*FET*_ < 10^−26^), and a *<*10% excess of genes that are differentially expressed in both head and gonad between males and females (*P*_*FET*_ = 0.036). We also get the same result when analyzing only two replicates of each sample: a significant excess of genes is differentially expressed in both head and testis between *Y*^*M*^ and *III*^*M*^ males (*P*_*FET*_ < 10^−4^), but not between males and females in both head and gonad (*P*_*FET*_ > 0.3). These results suggest that there are many genes under common regulatory control by the *M*-bearing chromosome in both male head and testis, but there is not the same degree of sex-specific regulation in common between head and gonad.

### 3.3. Genes that are differentially expressed between Y^M^ and III^M^ males are more likely to have male-biased expression

Genes whose expression is significantly higher in males than females are said to have “male-biased” expression, and genes that are up-regulated in females have “female-biased” expression (Ellegren and Parsch, 2007). We found that genes with male-biased expression in head are more likely to be differentially expressed between *Y*^*M*^ and *III*^*M*^ male heads than genes with either female-biased or unbiased expression (Figure 3A). Similarly, genes that are up-regulated in testis relative to ovary (“testis-biased”) are more likely to be differentially expressed between *Y*^*M*^ and *III*^*M*^ testes than genes with “ovary-biased” or unbiased expression in gonad (Figure 3B). We repeated this analysis using two replicates of each sample, and we consistently observe that genes with male-biased expression in head or gonad are more likely to be differentially expressed between *Y*^*M*^ and *III*^*M*^ males (Figures S4 and S5).

**Figure 3:**
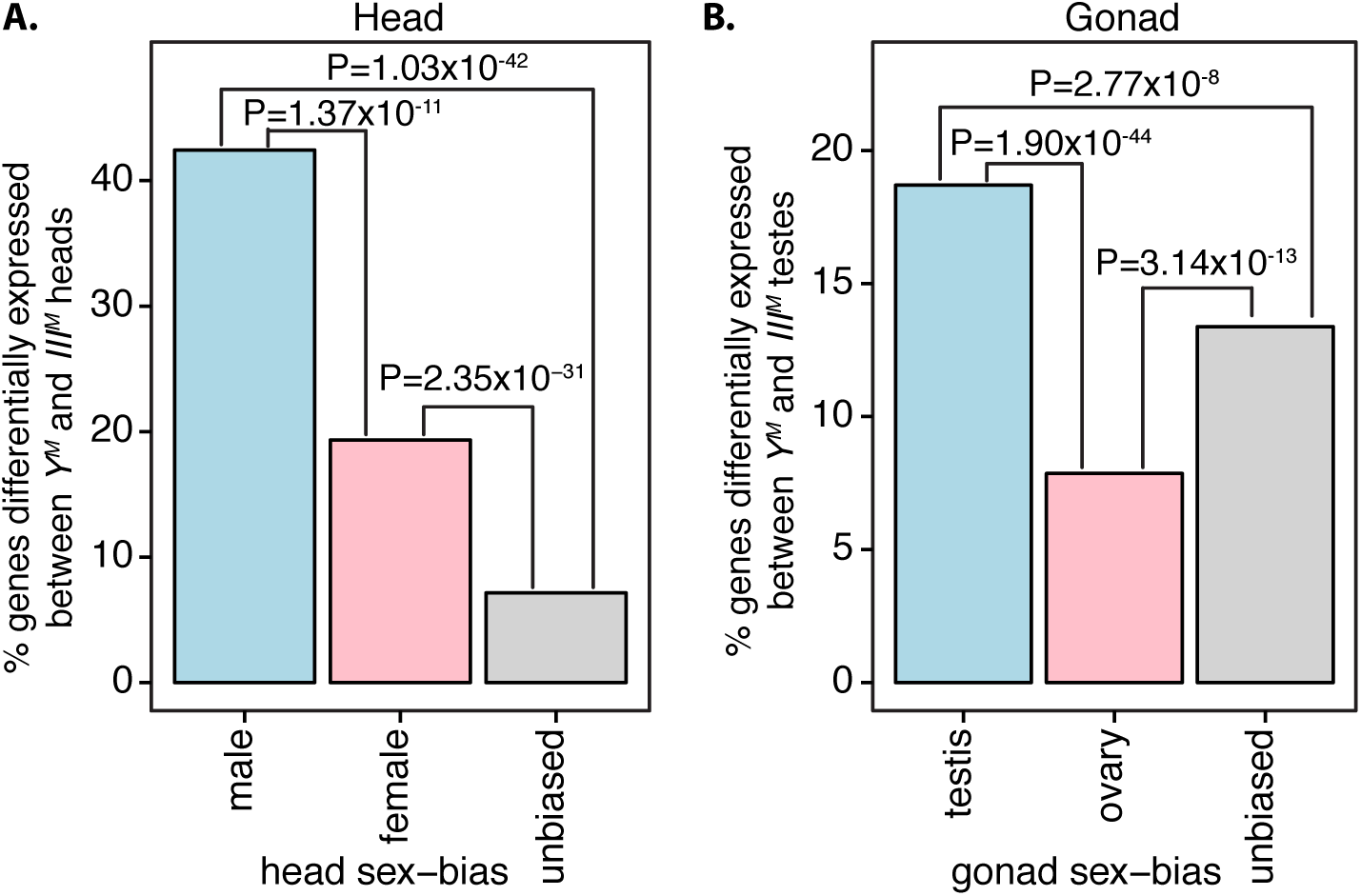
Sex-biased expression of genes differentially expressed between Y^M^ and III^M^ males. Bar graphs indicate the percentage of genes with male-biased (blue), female-biased (pink), or unbiased (gray) expression in either (A) head or (B) gonad that are differentially expressed between Y^M^ and III^M^ males in either (A) head or (B) testis. *P*-values are for Fisher’s exact test between groups.

### 3.4. Functional annotations of genes that are differentially expressed between Y^M^ and III^M^ males

We tested for GO categories that are over-represented amongst genes with sex-biased expression that are differentially expressed between *Y*^*M*^ and *III*^*M*^ males (Supplementary Data). We found that nearly half (49.7%) of genes that are differentially expressed between *Y*^*M*^ and *III*^*M*^ male heads are annotated with the functional category “catalytic activity”, whereas only 43% of genes not differentially expressed have that GO annotation (*P*_*FET*_ < 0.05 corrected for multiple tests). Over 10% of the genes with the catalytic activity annotation that are up-regulated in *III*^*M*^ male head are predicted to encode proteins involved in a metabolic process, including metabolism of organic acids, amino acids, and lipids. Among those genes, 15 are annotated as cytochrome P450 (CYP450) genes, and four of those also have male-biased expression in head (Table S6). CYP450s collectively carry out a wide range of chemical reactions including metabolism of endogenous (e.g., steroid hormones) and exogenous (e.g., xenobiotics) compounds (Scott, 2008). All 15 differentially expressed CYP450s are up-regulated in *III*^*M*^ males, and no CYP450 genes are up-regulated in *Y*^*M*^ males. Five of the CYP450s are on scaffolds that we assign to the third chromosome (Table S6), suggesting that *cis* regulatory sequences controlling the expression of CYP450s have diverged between *III*^*M*^ and the standard third chromosome. However, five of the CYP450s can be assigned to other autosomes (the remaining 5 cannot be assigned to a chromosome), demonstrating that divergence of *trans* factors between *III*^*M*^ and the standard third chromosome are also responsible for differential expression of CYP450s between *Y*^*M*^ and *III*^*M*^ males. The 15 CYP450s represent a range of different clans (2, 3 and 4) and families (4, 28, 304, 313, 438 and 3073) (Scott et al., 2014). However, an excess of CYP450s from clan 4 are up-regulated in *III*^*M*^ male head (*χ*^2^ = 4.19, *P* = 0.041), and thus overexpression of CYP450s is not random.

Genes that are annotated as encoding proteins located in extracellular regions are overrepresented amongst genes with testis-biased expression (15.0% of genes with testis-biased expression; 9.9% of genes not differentially expressed between testis and ovary; *P*_*FET*_ < 10^−3^ corrected for multiple tests) and amongst genes that are differentially expressed between *Y*^*M*^ and *III*^*M*^ testes (13.9% of differentially expressed genes; 8.1% of non-differentially expressed genes; *P*_*FET*_ < 10^−4^ corrected for multiple tests). In addition, 3.1% of the genes differentially expressed between *Y*^*M*^ and *III*^*M*^ testes are predicted to encode carbohydrate binding proteins (compared to 1.4% of non-differentially expressed genes; *P*_*FET*_ < 0.05 corrected for multiple tests), and 7.2% of differentially expressed genes are predicted to encode structural molecules (compared to 3.7% of non-differentially expressed genes; *P*_*FET*_ < 10^−3^ corrected for multiple tests). Three of those structural molecules are predicted to be *β*-tubulin proteins encoded by genes that are up-regulated in *Y*^*M*^ testes relative to *III*^*M*^, and two of those genes also have testis-biased expression. We tested for differential expression of the two of the *β*tubulin genes with testis-biased expression using qPCR (Figure S6; Supplementary Data). Only one of the two (XM 005187368) was up-regulated in *Y*^*M*^ testis when assayed with qPCR (*P <* 10^−4^), whereas the other gene (XM 005175742) was not (*P* = 0.653) possibly because of high variance in the *Y*^*M*^ measurement (Figure S6). The *D. melanogaster* genome encodes a testis-specific *β*-tubulin paralog that is essential for spermatogenesis (Kemphues et al., 1982; Hoyle and Raff, 1990), suggesting that the *β*-tubulin gene that is up-regulated in *Y*^*M*^ testis may be important for sperm development.

Four genes that are differentially expressed between *Y*^*M*^ and *III*^*M*^ testes are homologs of the *D. melanogaster Y*-linked fertility factors *kl-2*, *kl-3*, and *kl-5* (Goldstein et al., 1982; Gepner and Hays, 1993; Carvalho et al., 2000, 2001). These proteins encode components of the dynein heavy chain, which is necessary for flagellar activity of sperm. All four of the predicted dynein heavy chain genes that are differentially expressed between *Y*^*M*^ and *III*^*M*^ testes are autosomal in house fly. Three of these genes have testis-biased expression— two of those are up-regulated in *III*^*M*^ testis (XM 005175130, XM 005176585), while the third is up-regulated in *Y*^*M*^ testis (XM 005184828). The fourth gene (XM 005184771) is up-regulated in *III*^*M*^ testis, but it is not differentially expressed between testis and ovary. Using qPCR, we validated that XM 005184828 is up-regulated in *Y*^*M*^ testis (*P <* 10^−4^) and XM 005176585 is up-regulated in *III*^*M*^ testis (*P <* 0.01) (Figure S6; Supplementary Data). Two additional genes encoding components of other dynein chains have testis-biased expression and are up-regulated in *III*^*M*^ testis relative to *Y*^*M*^ testis.

Finally, there are numerous predicted RNAs in the house fly genome annotation that have no identifiable homology to any known RNAs or proteins (Scott et al., 2014). We identified six of these uncharacterized RNAs that both have testis-biased expression and are differentially expressed between *Y*^*M*^ and *III*^*M*^ testes. XR 225504, XR 225520, and XR 225639 are up-regulated in *Y*^*M*^ testes, and XR 225442, XR 225497, and XR 225737 are up-regulated in *III*^*M*^ testes. These genes are annotated as encoding non-coding RNAs (ncRNAs), and we were unable to detect long open reading frames in the transcripts. It is possible that these ncRNAs are responsible for the regulation of gene expression in testis, and differential expression of these ncRNAs between *Y*^*M*^ and *III*^*M*^ testis could be responsible for the differential expression of other genes between *Y*^*M*^ and *III*^*M*^ males.

## 4. Discussion

### 4.1. Differential expression between Y^M^ and III^M^ males is driven by both cis and trans effects

We compared gene expression in head and testis between *Y*^*M*^ and *III*^*M*^ males. *Y*^*M*^ males are homozygous for a standard third chromosome that does not have *M*, whereas *III*^*M*^ males are heterozygous for a *III*^*M*^ chromosome and a standard third chromosome (Figure 1). Differences in the expression levels of autosomal genes between *Y*^*M*^ and *III*^*M*^ males could be the result of: 1) divergence of *cis*-regulatory sequences between the *III*^*M*^ and standard third chromosomes that affect the expression of genes on the third chromosome; 2) divergence of *trans*-factors between *III*^*M*^ and the standard third chromosome that differentially regulate gene expression throughout the genome; 3) downstream effects of the first two processes that lead to further differential expression.

The two strains also differ in the genotype of their sex chromosomes; *Y*^*M*^ males are *XY*, whereas *III*^*M*^ males are *XX* (Figure 1). The house fly *Y* chromosome is highly heterochromatic and does not harbor any known genes other than *M* (Boyes et al., 1964; Hediger et al., 1998; Dübendorfer et al., 2002). It is clear that the *Y* chromosome does not contain any genes necessary for male fertility or viability because *XX* males are fertile and viable. The *X* chromosome is also highly heterochromatic and probably homologous to the *Drosophila* dot chromosome (Hediger et al., 1998; Vicoso and Bachtrog, 2013, 2015). The heterochromatic *Drosophila Y* chromosome can affect the expression of autosomal genes (Lemos et al., 2008, 2010; Sackton et al., 2011; Zhou et al., 2012), suggesting that the house fly *X* and *Y* chromosomes could have *trans* regulatory effects on autosomal gene expression.

A higher fraction of third chromosome genes are differentially expressed between *Y*^*M*^ and *III*^*M*^ house fly males than genes on any other autosome (Figure 2A). Therefore, divergence of *cis*-regulatory sequences between *III*^*M*^ and the standard third chromosome are at least partially responsible for the expression differences between *Y*^*M*^ and *III*^*M*^ males. However, ˜70% of the genes differentially expressed between *Y*^*M*^ and *III*^*M*^ males map to one of the other four autosomes, suggesting that the majority of expression differences are the result of *trans* effects of the *X*, *Y*, and third chromosomes along with further downstream effects.

### 4.2. Reproductive and male phenotypes are more likely to be affected by M variation

Reproductive traits are more sexually dimorphic than non-reproductive traits, and reproductive traits also tend to evolve faster, possibly as a result of sexual selection (Eberhard, 1985). A similar faster evolution of gene expression in reproductive tissues has been observed across many taxa (Khaitovich et al., 2005; Zhang et al., 2007; Brawand et al., 2011), and increased variation within species for sex-biased gene expression often accompanies elevated expression divergence (Meiklejohn et al., 2003; Ayroles et al., 2009). Consistent with these patterns, more genes are differentially expressed between *Y*^*M*^ and *III*^*M*^ males in testis than head (Table 1, Figure S2). Somatic and germline SD in house fly are under the same genetic control (Hilfiker-Kleiner et al., 1994), so the exaggerated differences in expression between *Y*^*M*^ and *III*^*M*^ testes relative to heads cannot be attributed to differences in the SD pathway between gonad and head. We also find that genes with male-biased expression are more likely to be differentially expressed between *Y*^*M*^ and *III*^*M*^ males (Figure 3). Genes with male-biased expression are more likely to perform sex-specific functions (Connallon and Clark, 2011), suggesting that genes that are differentially expressed between *Y*^*M*^ and *III*^*M*^ males disproportionately affect male phenotypes.

### 4.3. Evaluating the role of sex-specific selection in MSD turnover

Many models of SD evolution predict that a new MSD locus will invade a population if it is genetically linked to an allele with a beneficial, sexually selected, or sexually antagonistic fitness effect (Charlesworth and Charlesworth, 1980; Rice, 1987; Charlesworth, 1991, 1996; Rice, 1996; van Doorn and Kirkpatrick, 2007, 2010). Alternatively, evolutionary turnover of MSD loci could be the result of neutral drift in a highly labile system (van Doorn, 2014).

Our results are consistent with a model in which the *III*^*M*^ chromosome invaded because it harbors alleles with male-specific beneficial effects. First, the expression of genes that are likely to perform male-specific functions—especially in male fertility—are more likely to be affected by the *III*^*M*^ chromosome (Table 1; Figure 3), and those male-specific phenotypic differences could have been targets of sex-specific selection pressures. However, as mentioned above, the expression of male-biased genes is more variable than other genes even in species without multifactorial SD systems (Meiklejohn et al., 2003; Ayroles et al., 2009). Additional experiments in which a non-*M*-bearing chromosome is placed on a common genetic background are therefore necessary to further test the hypothesis that the *M*-bearing chromosome disproportionately affects male-biased gene expression.

We also found that *III*^*M*^ heads have a masculinized expression profile relative to *Y*^*M*^ heads, suggesting that the male-limited transmission of the *III*^*M*^ chromosome favored the accumulation of alleles with male-beneficial fitness effects (Rice, 1984). Previous work found that *III*^*M*^ males outperformed *Y*^*M*^ males in multiple laboratory fitness assays (Hamm et al., 2009), providing additional support for the accumulation of male-beneficial alleles on the *III*^*M*^ chromosome. However, despite the apparent selective advantage of the *III*^*M*^ chromosome, it surprisingly does not appear to be expanding rapidly (Hamm et al., 2015), suggesting that the fitness benefits of *III*^*M*^ may be environment-specific (Feldmeyer et al., 2008).

Our data do not allow us to distinguish between two possible orders of events in the invasion of the *III*^*M*^ chromosome. In the first scenario, male-beneficial alleles on the third chromosome could have driven the initial invasion of *III*^*M*^ (van Doorn and Kirkpatrick, 2007). In the second scenario, beneficial alleles could have accumulated on the *III*^*M*^ chromosome after it acquired an *M*-locus because male-limited inheritance promotes the fixation of male-beneficial alleles (Rice, 1984, 1987). These scenarios are not mutually exclusive. Regardless of the sequence of events, we have provided evidence that the house fly multifactorial male-determining system is associated with phenotypic differences that likely have male-specific fitness effects, which could explain the invasion of the *III*^*M*^ chromosome under sexual or sexually antagonistic selection.

We also found that 14.8% of genes that are up-regulated in *III*^*M*^ male heads have male-biased expression, whereas *<*2% of genes that are up-regulated in *Y*^*M*^ male heads have male-biased expression (*P*_*FET*_ < 10^−10^). We observe the same excess of male-biased genes up-regulated in *III*^*M*^ male heads when we only use two replicates of each strain and sex to test for differential expression (*P*_*FET*_ < 0.05 using most combinations of two replicates). This suggests that *III*^*M*^ male heads have a “masculinized” expression profile relative to *Y*^*M*^ heads.

## 5. Acknowledgements

Cheryl Leichter, Naveen Galla, and Daniel Chazen assisted with the creation of the CSaY strain, Amanda Manfredo prepared the mRNA-Seq libraries, and Christopher Gonzales performed the qPCR validation. This work benefited from discussions with members of the Clark lab. We were supported by multistate project S-1030 to JGS, NIH grant R01GM64590 to AGC and A. Bernardo Carvalho, and start-up funds from the University of Houston to RPM. The funders had no role in study design, data collection and analysis, or preparation of the manuscript.

## Supplementary Tables

**Table S1:**
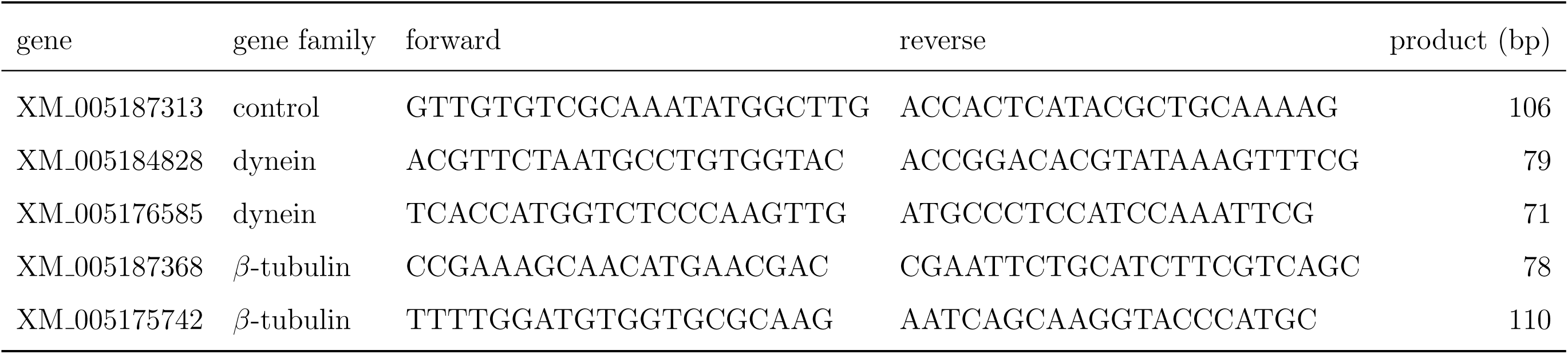
qPCR primers to test for differential expression.

**Table S2:** Differential expression between strains and sexes using the same genes in gonad and head tissue samples. Same as Table 1 except only genes that are expressed in both head and gonad are counted.

| Tissue | comparison | # diff |  | # non-diff | # genes | freq diff |
| --- | --- | --- | --- | --- | --- | --- |
| | | $Y^M$ | $III^M$ | | | |
| male head | $Y^M$ vs. $III^M$ | 260 | 426 | 7628 | 8314 | 0.083 |
| testis | $Y^M$ vs. $III^M$ | 587 | 386 | 7341 | 8314 | 0.117 |
|  |  | male | female |  |  |  |
| head | male vs. female | 143 | 931 | 7522 | 8596 | 0.125 |
| gonad | testis vs. ovary | 2386 | 3989 | 2221 | 8596 | 0.742 |

**Table S3:** Differential expression between strains and sexes using only two replicates. Same as Table 1 except only two biological replicates of male samples were used in the analysis to control for sample-size effects. In the Y^M^ vs. III^M^ comparisons, we present the average of the number of genes differentially expressed between 1) the two Y^M^ strains that are best correlated and the two III^M^ strains that are best correlated; the two Y^M^ strains that are worst correlated and the two III^M^ strains that are worst correlated; 3) a third pair of Y^M^ strains and a third pair of III^M^ strains. Correlations can be seen in Figure S1. In the male vs. female and testis vs. ovary comparisons, we present the average of the number of genes differentially expressed between 1) the first two Y^M^ and III^M^ replicates and the second two female replicates; 2) the second two Y^M^ and III^M^ replicates and the second two female replicates; 3) the third two Y^M^ and III^M^ replicates and the third two female replicates.

| Tissue | comparison | # diff |  | # non-diff | # genes | freq diff |
| --- | --- | --- | --- | --- | --- | --- |
| | | $Y^M$ | $III^M$ | | | |
| male head | $Y^M$ vs. $III^M$ | 321 | 316 | 8829 | 9465 | 0.067 |
| testis | $Y^M$ vs. $III^M$ | 602 | 482 | 9285 | 10365 | 0.104 |
|  |  | male | female |  |  |  |
| head | male vs. female | 232 | 170 | 9891 | 10293 | 0.039 |
| gonad | testis vs. ovary | 3549 | 3308 | 3616 | 10473 | 0.655 |

**Table S4:**
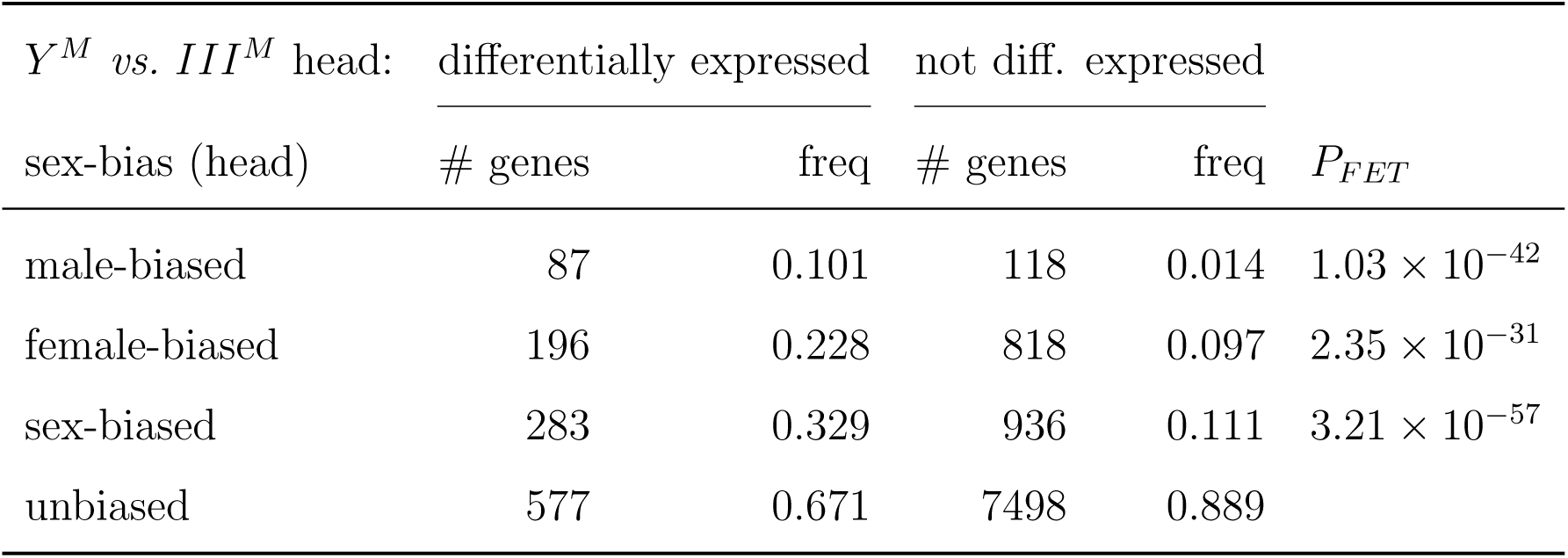
Sex-biased expression in head of genes that are differentially expressed between Y^M^ and III^M^ male heads. Sex-biased expression is based on comparisons of male and female heads. The number of genes (# genes) that are differentially expressed between strains and non-differentially expressed is reported for each sex-bias category. We also report the frequency of genes within the differential expression class that are in each sex-bias category (freq). A *P*-value is given for Fisher’s exact test comparing differentially and non-differentially expressed genes in each sex-bias category with genes that have unbiased expression (P_F ET_).

**Table S5:** Sex-biased expression in gonad of genes that are differentially expressed between Y^M^ and III^M^ testes. Sex-biased expression is based on comparisons of testis and ovary. The number of genes (# genes) that are differentially expressed between strains and non-differentially expressed is reported for each sex-bias category. We also report the frequency of genes within the differential expression class that are in each sexbias category (freq). A *P*-value is given for Fisher’s exact test comparing differentially and non-differentially expressed genes in each sex-bias category with genes that have unbiased expression (P_F ET_).

| $Y^M$ vs. $III^M$ testis: | differentially expressed | | not diff. expressed | | $P$-value |
| --- | --- | --- | --- | --- | --- |
|  | # genes | freq | # genes | freq |  |
| sex-bias (gonad) |  |  |  |  |  |
| testis | 629 | 0.481 | 2735 | 0.309 | $2.77 \times 10^{-8}$ |
| ovary | 325 | 0.248 | 3810 | 0.431 | $3.14 \times 10^{-13}$ |
| sex-biased | 954 | 0.729 | 6545 | 0.740 | 0.381 |
| unbiased | 355 | 0.271 | 2298 | 0.260 |  |

**Table S6:**
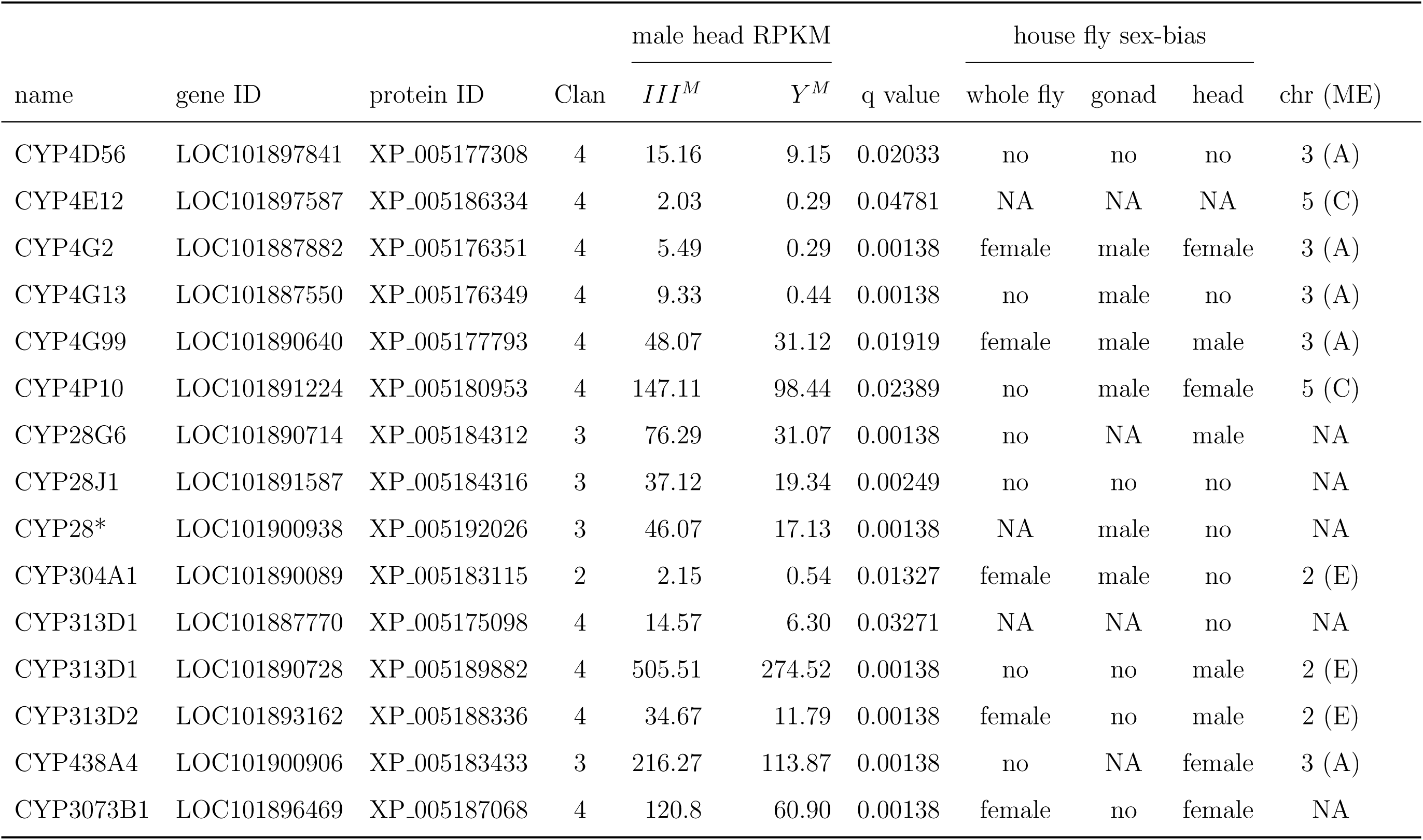
CYP450 genes that are differentially expressed between III^M^ and Y^M^ male heads. The asterisk (*) indicates a gene with an incomplete sequence in the assembly. The q value is an FDR-corrected P value comparing expression in III^M^ and Y^M^ heads. Sex-biased expression (male, female, and no) is given for whole fly measurements (Scott et al., 2014), gonad, and head. Genes for which expression level was too low to test for sex-biased expression are given as NA. House fly chromosome arm assignments (and Muller element homologies) are provided for genes that are on scaffolds where the majority of 1:1 D. melanogaster homologs are on a single Muller element (NA is listed for genes whose scaffolds could not be assigned to a chromosome arm).

## Supplementary Figures

**Figure S1:**
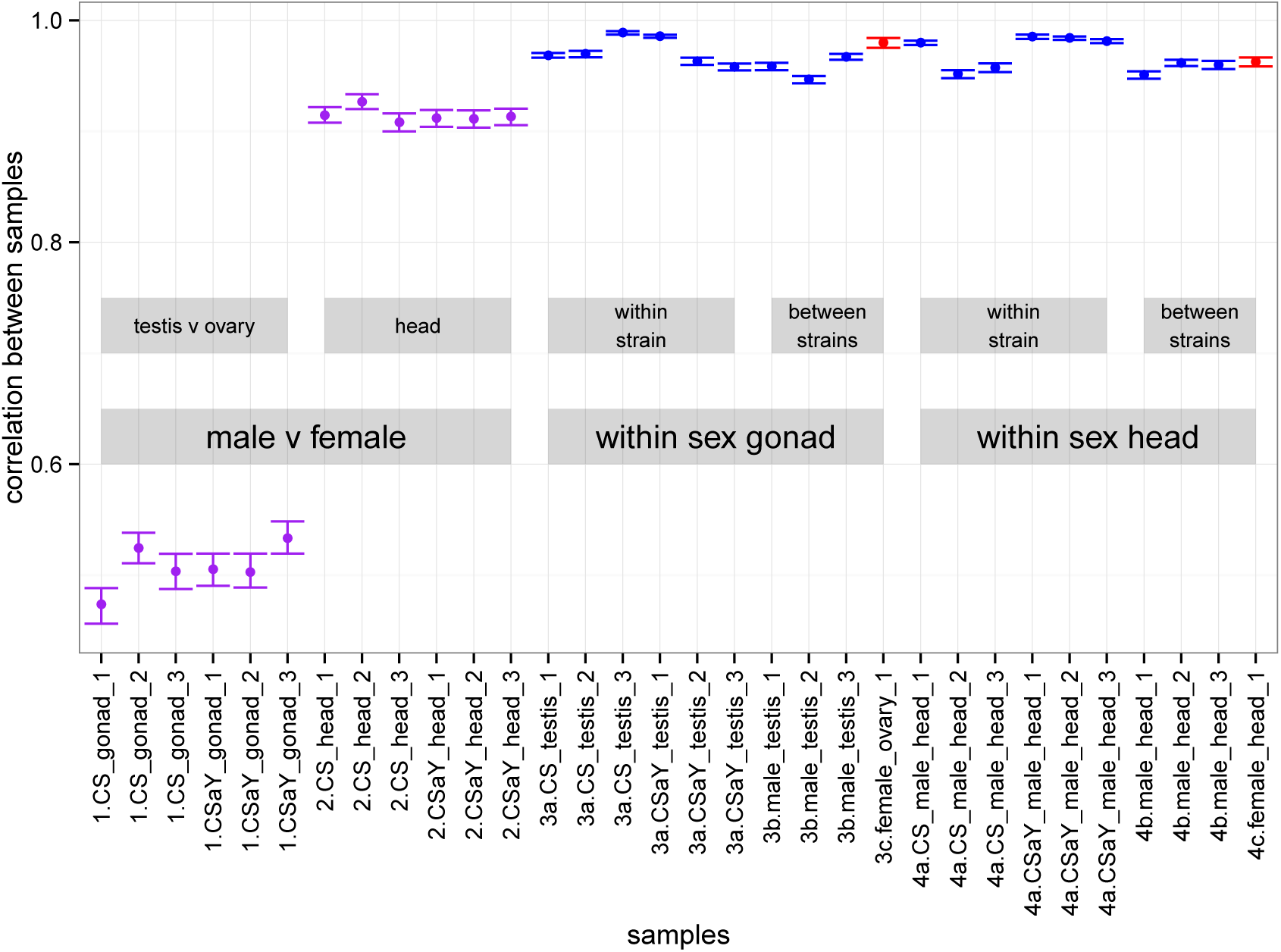
Correlations of expression levels of genes between samples. Point estimates of Spearman’s correlation coefficient (ρ) of gene expression between samples are plotted, along with the 95% confidence intervals (CIs) of the correlations. CIs were estimated by bootstrapping the data 1000 times. Comparisons between male and female samples are colored purple, comparisons between male samples are colored blue, and comparisons between female samples are colored red.

**Figure S2:**
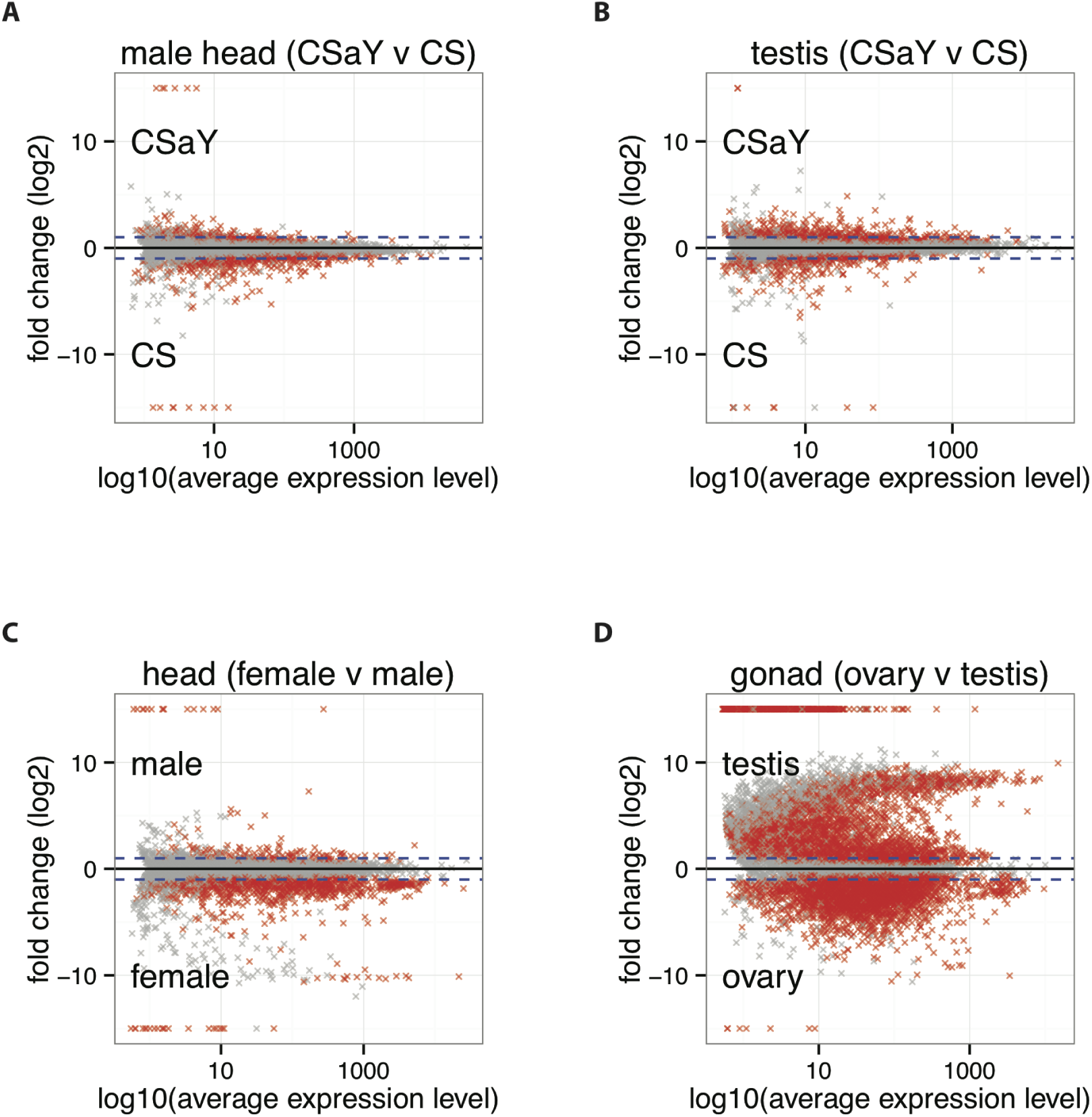
MA plots for comparisons of expression between (A) male heads from the CSaY (Y^M^) and CS (III^M^) strains, (B) testes from the CSaY and CS strains, (C) heads from females and males, and (D) testes and ovaries. The fold change (*log_2_* expression ratio) is plotted against *log_10_* of the average expression in the two samples being compared. Gray points represent genes whose expression does not significantly differ between strains or sexes (darker gray indicates a higher density of points), and red points represent genes whose expression does differ significantly. Points at the top or bottom of the plot (*log_2_* fold change −15 or 15) are genes with an expression value of zero in one strain or sex. Only genes that were tested for significantly differential expression are plotted. The dashed blue lines are the 2-fold cutoffs.

**Figure S3:**
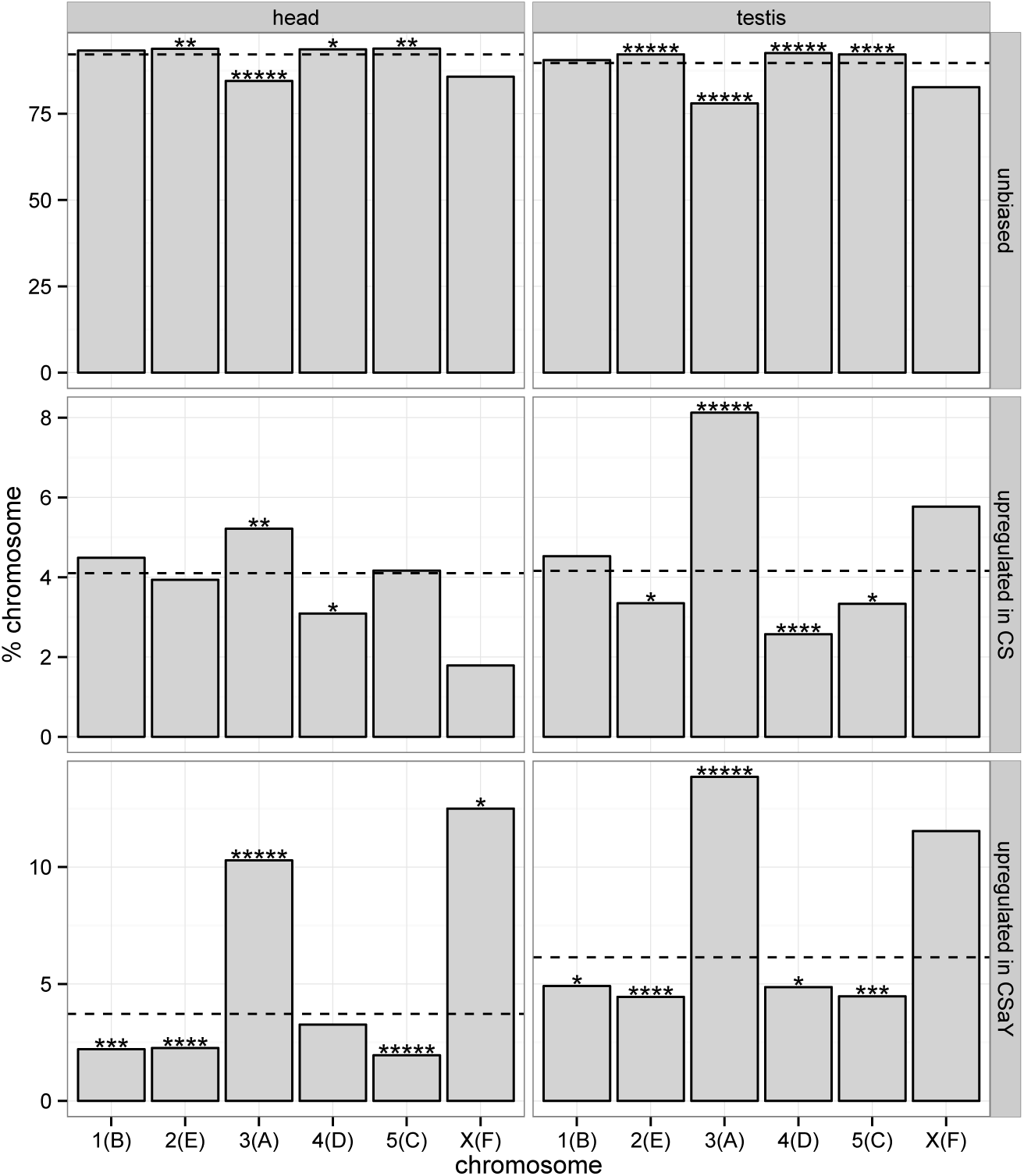
Chromosomal locations of genes that are differentially expressed between Y^M^ and III^M^ males. Same as Figure 2 except that differentially expressed genes are separated into those that are up-regulated in III^M^ (CS) males (middle), or up-regulated in Y^M^ (CSaY) males (bottom).

**Figure S4:**
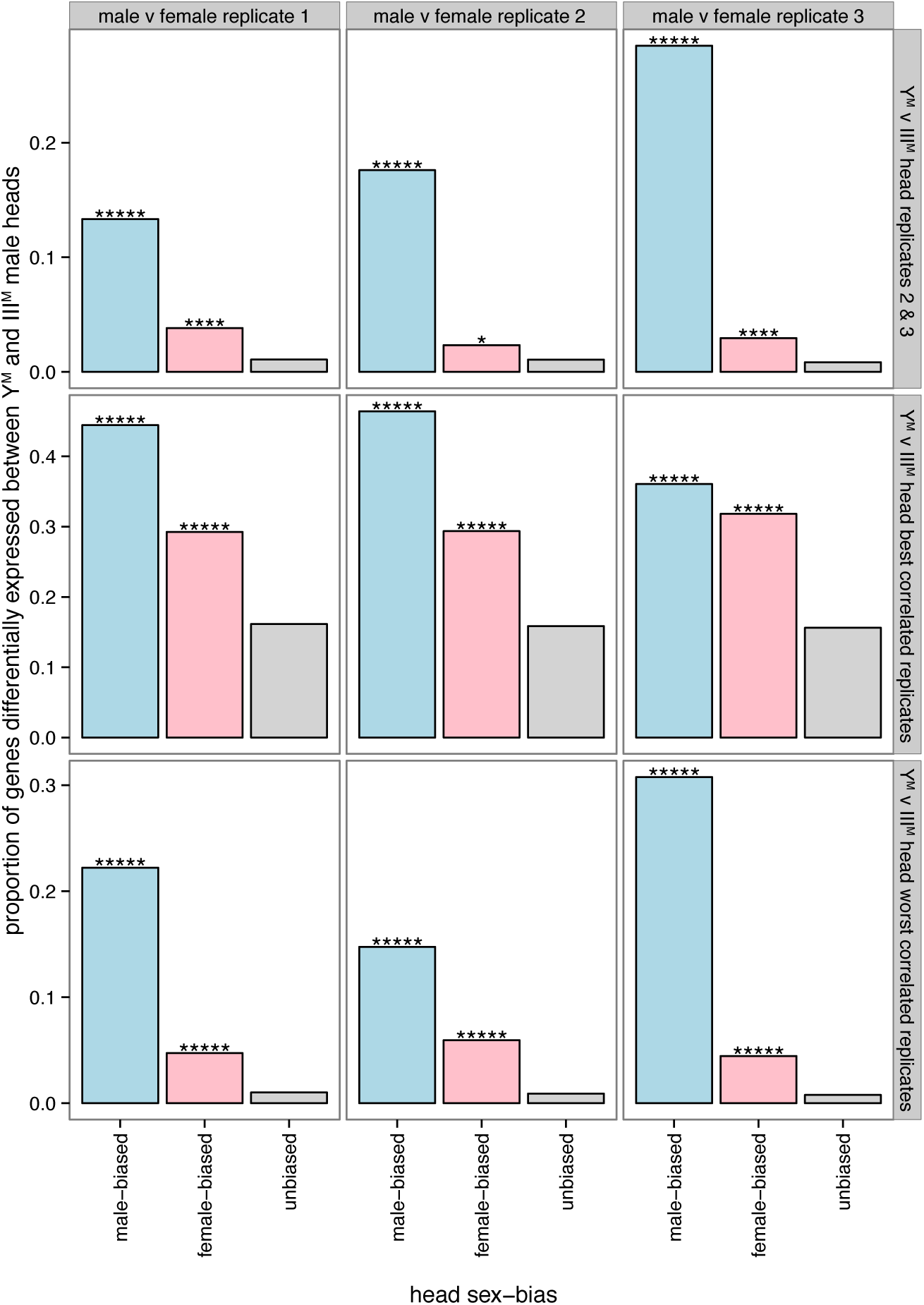
Sex-biased expression of genes differentially expressed between Y^M^ and III^M^ heads. Same as Figure 3 except only two replicates of each strain were used to test for differential expression between Y^M^ and III^M^ male heads and between male and female heads. Three different subsets of the data were used for the male-female comparison (listed along the top of the figure). Three different subsets were used in the Y^M^ vs. III^M^ comparisons: 1) a random pair of Y^M^ strains and a pair of III^M^ strains; 2) the two Y^M^ strains that are best correlated and the two III^M^ strains that are best correlated; 3) the two Y^M^ strains that are worst correlated and the two III^M^ strains that are worst correlated. Asterisks indicate significant differences between the proportion of genes with sex-biased and unbiased expression determined using Fisher’s exact test (* P < 0.05, ** P < 0.005, *** P < 0.0005, **** P < 0.00005, ***** P < 0.000005).

**Figure S5:**
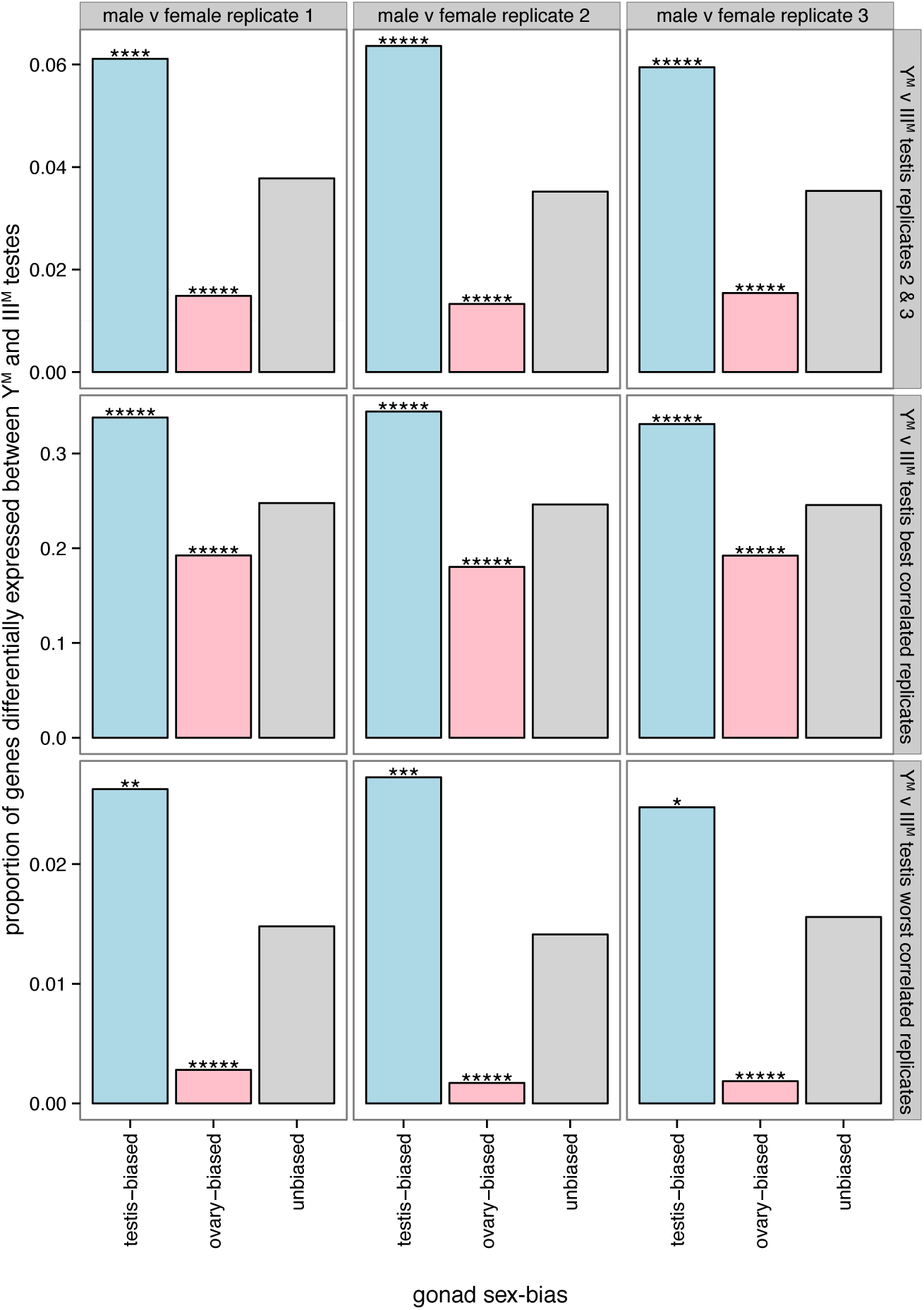
Sex-biased expression of genes differentially expressed between Y^M^ and III^M^ testis. Same as Figure 3 except only two replicates of each strain were used to test for differential expression between Y^M^ and III^M^ testes and between testis and ovary. Three different subsets of the data were used for the male-female comparison (listed along the top of the figure). Three different subsets were used in the Y^M^ vs. III^M^ comparisons: 1) a random pair of Y^M^ strains and a pair of III^M^ strains; 2) the two Y^M^ strains that are best correlated and the two III^M^ strains that are best correlated; 3) the two Y^M^ strains that are worst correlated and the two III^M^ strains that are worst correlated. Asterisks indicate significant differences between the proportion of genes with sex-biased and unbiased expression determined using Fisher’s exact test (* P < 0.05, ** P < 0.005, *** P < 0.0005, **** P < 0.00005, ***** P < 0.000005).

**Figure S6:**
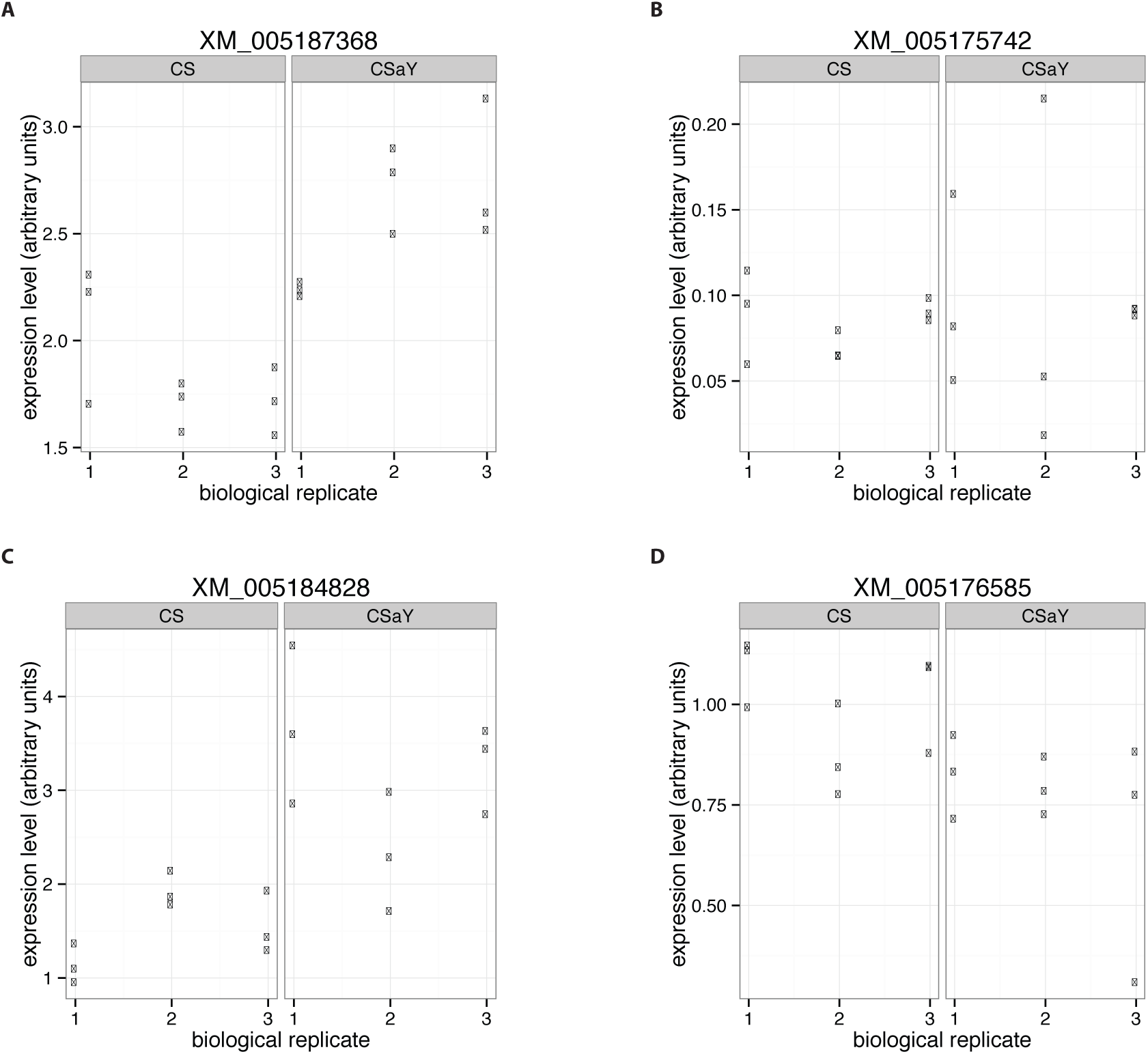
Estimates of expression level from the qPCR assay of four genes are plotted. Three technical replicates of three biological replicates were measured. Results of the statistical analyses are presented in the main body of the manuscript.

## Supplementary Data

### mRNA-Seq results

Column information:

1. **gene** - NCBI gene symbol
2. **protein id** - NCBI protein identifier
3. **locus** - Coordinates in genomic scaffold
4. **orthocat** - Orthology category:
  - sco: single-copy ortholog
  - mto1: multiple house fly genes orthologous to 1 *D. melanogaster* gene
  - 1tom: 1 house fly gene orthologous to many *D. melanogaster* genes
  - mtom: multiple house fly genes homologous to multiply *D. melanogaster* genes
  - lr: lineage-restricted house fly gene without any identified *D. melanogaster* orthologs
5. **og** - Orthology group (gene family)
6. **dmel** - FlyBase identifier of *D. melanogaster* orthologs
7. **Dmel_ME** - Muller element location of *D. melanogaster* ortholog (only if sco)
8. **Musca_chr** - House fly chromosome homolog of Muller element
9. **scaff_ME** - Muller element assignment of house fly scaffold (based on majority rule of genes on scaffold)
10. **Musca_scaff** - House fly chromosome based on scaff ME
11. **CS_MaleHead_FPKM** - Expression level in CS (*III*^*M*^) male head
12. **CSaY_MaleHead_FPKM** - Expression level in CSaY (*Y*^*M*^) male head
13. **MaleHead_TestStat** - Test statistic comparing CS and CSaY male head expression levels
14. **MaleHead_qval** - FDR-corrected *P*-value of test statistic comparing CS and CSaY male head expression levels
15. **CS_testis_FPKM** - Expression level in CS (*III*^*M*^) testis
16. **CSaY_testis_FPKM** - Expression level in CSaY (*Y*^*M*^) testis
17. **testis_TestStat** - Test statistic comparing CS and CSaY testis expression levels
18. **testis_qval** - FDR-corrected *P*-value of test statistic comparing CS and CSaY testis expression levels
19. **CS_FemaleHead_FPKM** - Expression level in CS female head
20. **CSaY_FemaleHead_FPKM** - Expression level in CSaY female head
21. **FemaleHead_TestStat** - Test statistic comparing CS and CSaY female head expression levels
22. **FemaleHead_qval** - FDR-corrected *P*-value of test statistic comparing CS and CSaY female head expression levels
23. **CS_ovary_FPKM** - Expression level in CS ovary
24. **CSaY_ovary_FPKM** - Expression level in CSaY ovary
25. **ovary_TestStat** - Test statistic comparing CS and CSaY ovary expression levels
26. **ovary_qval** - FDR-corrected *P*-value of test statistic comparing CS and CSaY ovary expression levels
27. **female_head_FPKM** - Expression level in female head
28. **male_head_FPKM** - Expression level in CS male head
29. **head_TestStat** - Test statistic comparing female and male head expression levels
30. **head_qval** - FDR-corrected *P*-value of test statistic comparing female and male head expression levels
31. **ovary_FPKM** - Expression level in ovary
32. **testis_FPKM** - Expression level in testis
33. **gonad_TestStat** - Test statistic comparing ovary and testis expression levels
34. **gonad_qval** - FDR-corrected *P*-value of test statistic comparing ovary and testis expression levels

### Gene ontology tests

File information (test vs reference comparisons):

1. head_female-biased_GO.txt - GO categories that are enriched in genes with female-biased expression in head (foreground) vs genes with non-sex-biased expression in head
2. head_male-biased_GO.txt - GO categories that are enriched in genes with male-biased expression in head vs genes with non-sex-biased expression in head
3. male_head_diff_GO.txt - GO categories that are enriched in genes that are differentially expressed between CS and CSaY male heads vs genes that are not differentially expressed between CS and CsaY male heads
4. male_testis_diff_GO.txt - GO categories that are enriched in genes that are differentially expressed between CS and CSaY testes vs genes that are not differentially expressed between CS and CsaY testes
5. ovary-biased_unbiased_GO.txt - GO categories that are enriched in genes with ovary-biased expression vs genes with non-sex-biased expression in gonad
6. testis-biased_unbiased_GO.txt - GO categories that are enriched in genes with testis-biased expression vs genes with non-sex-biased expression in gonad

Column information:

1. GO-ID - Gene ontology ID
2. Term - GO term
3. Category - Cellular Component (C), Biological Process (P), or Molecular Function (F)
4. FDR - False discovery corrected *P*-value for enrichment
5. *P*-value - Fisher’s exact test comparing #Test, #Ref, #notAnnotTest, #notAnnotRef
6. #Test - Count of genes with ontology annotation that are differentially expressed
7. #Ref - Count of genes with ontology annotation that are not differentially expressed
8. #notAnnotTest - Count of genes without ontology annotation that are differentially expressed
9. #notAnnotRef - Count of genes without ontology annotation that are not differentially expressed
10. Over/Under - Term is enriched (over) or depleted (under) in test data set

### qPCR results

Raw qPCR data are presented for each of the four genes assayed (one file per gene). Each table has three columns: 1) gene name (XM 005187313 is the control gene, and the other gene is being tested for differential expression); 2) replicate (whether the data are from one of the serial dilutions used to calculate the standard curve or a biological replicate of the two strains being tested); 3) CT value of the technical replicate.

